# Glucose-dependent activation, activity, and deactivation of beta cell networks in acute mouse pancreas tissue slices

**DOI:** 10.1101/2020.03.11.986893

**Authors:** Andraž Stožer, Maša Skelin Klemen, Marko Gosak, Lidija Križančić Bombek, Viljem Pohorec, Marjan Slak Rupnik, Jurij Dolenšek

## Abstract

Many details of glucose-stimulated intracellular calcium changes in beta cells during activation, activity and deactivation, as well as their concentration-dependence, remain to be described. Classical physiological experiments indicated that in islets functional differences between individual cells are largely attenuated, but recent findings suggest considerable intercellular heterogeneity, with some cells possibly coordinating the collective responses. To address the above with an emphasis on heterogeneity and describing the relations between classical physiological and functional network properties, we performed functional multicellular calcium imaging in mouse pancreas tissue slices over a wide range of glucose concentrations. During activation, delays to activation of cells and first responder-any cell delays shortened, and the sizes of simultaneously responding clusters increased with increasing glucose. Exactly the opposite characterized deactivation. The frequency of fast calcium oscillations during activity increased with increasing glucose up to 12 mM glucose, beyond which oscillation duration became longer, resulting in a homogenous increase in active time. In terms of functional connectivity, islets progressed from a very segregated network to a single large functional unit with increasing glucose. A comparison between classical physiological and network parameters revealed that the first-responders during activation had longer active times during plateau and the most active cells during the plateau tended to deactivate later. Cells with the most functional connections tended to activate sooner, have longer active times, and deactivate later. Our findings provide a common ground for recent differing views on beta cell heterogeneity and an important baseline for future studies of stimulus-secretion and intercellular coupling.

## 1. Introduction

Glucose is the main insulin secretagogue in beta cells. After entering beta cells through GLUT (131) or other transporters (117), glucose initiates cellular processes leading to transient activation of beta cells, followed by subsequent periodic oscillatory changes in membrane potential and intracellular calcium concentration ([Ca^2+^]_IC_, as well as insulin secretion (22, 23, 61, 72, 111, 138). The calcium ion (Ca2+) is the central secondary messenger, coupling stimulation by nutrients and many other neurohormonal stimuli with secretion (58, 73, 75, 131). Glucose-dependent oscillatory [Ca^2+^]_IC_ changes have been reported in beta cells *in vivo* in an exteriorized pancreas and in islets transplanted into the anterior eye chamber, *in situ* in acute pancreas tissue slices, and *in vitro* in enzymatically isolated and cultured islets and isolated cells (17, 19, 23, 26, 33, 54, 69, 134, 135). [Ca^2+^]_IC_ oscillations are the intermediate or central step in beta cell stimulus-secretion coupling. On the one hand, they are well synchronized with membrane potential oscillations or so-called bursts of electrical activity (47, 59, 125). On the other hand, they are the triggering signal for the fusion of insulin-containing vesicles with the plasma membrane, thus driving the periodic release of insulin (23, 60, 61). The extent of insulin secretion is further regulated by various additional signals in both [Ca^2+^]_IC_-dependent and [Ca^2+^]_IC_-independent manner (1, 29, 73, 131). Regardless of the pathways taken to induce and regulate insulin secretion, [Ca^2+^]_IC_ remains a convenient proxy for assessing both more proximal and more distal events in the stimulus-secretion coupling cascade. Compared with measurements of proximal membrane potential changes and distal insulin secretion, at present high frequency confocal functional multicellular [Ca^2+^]_IC_ imaging typically offers the best combination of the number of cells that can be analyzed simultaneously, spatial and temporal resolution, and total recording time (2, 43, 46, 66, 101, 139, 140, 146).

Beta cells within an islet typically display a biphasic [Ca^2+^]_IC_ response to physiological stimuli (23, 59, 67, 101, 110, 139, 140, 142). From a temporal perspective, this biphasic [Ca^2+^]_IC_ response is well in accordance with the biphasic insulin release observed *in vitro* in isolated islets (13, 60). An initial transient increase in [Ca^2+^]_IC_ is followed by a sustained plateau phase with superimposed fast [Ca^2+^]_IC_ oscillations. The frequency and duration of these fast [Ca^2+^]_IC_ oscillations closely resemble electrophysiologically measured bursts of mebrane potential depolarizations, as well as fast pulses of insulin secretion (11, 22, 24, 47, 59, 61, 125). During the transient phase, the initial activation of cells varies significantly in time and space. Beta cells activate in small clusters dispersed over an islet with no predictable pattern (47, 67, 142). During the plateau phase, [Ca^2+^]_IC_ oscillations are slightly phase-lagged due to intercellular depolarization, and [Ca^2+^]_IC_ waves repeatedly spread across islets with an average velocity of about 100 μm/s (10, 19, 30, 47, 80, 129, 139, 143). Simultaneous measurements of membrane potential and [Ca^2+^]_IC_ oscillations demonstrated a tight coupling of the two processes (16, 59, 125), with membrane potential depolarization preceding the [Ca^2+^]_IC_ increase by about 150 ms (47). After lowering glucose back to substimulatory concentration, beta cells in all types of preparations deactivate, and [Ca^2+^]_IC_ returns to the basal level, (39, 81, 101, 139, 140).

A substantial number of previous studies attempted to address the glucose-dependency of beta cell stimulation; however, a complete characterization of the complex relationship with stimulus intensity remains elusive. Both *ex vivo* (9, 20, 41, 57, 74, 97) and *in vivo* experiments (74) show that activity, assessed by measurements of membrane potential, [Ca^2+^]_IC_, or insulin secretion, progressively increases over a wide range of physiological and supraphysiological glucose concentrations. In dissociated beta cells in culture, glucose thresholds for single beta cell activation of [Ca^2+^]_IC_ differed substantially among individual beta cells, ranging from low to supraphysiological glucose concentrations. Increasing stimulus intensity progressively recruited additional beta cells into a secretory response (20, 85, 130), a phenomenon well explained by beta cell heterogeneity with respect to their sensitivity to glucose (50, 91, 142). However, beta cells in intact islets function in a coupled, collective way and such coupling significantly narrows the concentration range over which the whole islet activates. The half-maximal effective concentration of glucose (EC50) demonstrated for [Ca^2+^]_IC_ and membrane potential is about 7 mM, and the most significant increase in electrical activity in the range of about 6-8 mM (20, 40), indicating that additional mechanisms contribute to insulin release above the physiological glucose range. Among these, different sources of amplification, e.g., hormonal, neuronal, and metabolic, could also contribute to the elevated average [Ca^2+^]_IC_ driving increased insulin release in both uncoupled (85) and coupled beta cells (7, 70, 74). The most emphasized mechanism demonstrated already decades ago operates by changing the time that the cells spend in an active state during the plateau phase, by increasing either the frequency or the duration of membrane potential bursts. To the best of our knowledge, there are only a handful of studies that analyzed the glucose-dependence of active time in isolated islets (7, 37, 38, 76, 95, 104). All of these studies employed membrane potential measurements, which are inherently limited by a rather low number of successful impalements. Additionally, they usually predict that the electrical behavior in other cells is the same as in the observed cells and are therefore unable to quantify the heterogeneity of responses during activation and deactivation and compare it with the behavior during the plateau phase. The same holds true for complex network analyses of functional connectivity that require recordings from many cells at a time (66, 78, 79, 84, 101, 123, 140). Finally and most importantly, at least to some extent all of the studies employed sequential increasing or lowering of glucose during prolonged impalements and then analyzed short intervals in a given concentration. In such experimental setups, the effects of time *per se* or so-called memory effects cannot be separated from effects of different concentrations of glucose. In our present study, we tried to circumvent all of the above shortcomings.

It should be pointed out that the increase in [Ca^2+^]_IC_ probably exerts its main effect through increasing the number of fusion events of insulin-containing granules (97, 132). Recent mathematical modeling of experimental data corroborates this idea, demonstrating that both mobilization and priming of insulin granules are the main factors determining concentration-dependent insulin secretion (111, 138). In theory, the effects observed at supraphysiological glucose concentrations could also be due to mechanisms that do not operate at lower glucose concentrations. It is important to emphasize that although the physiological range of glucose in mice has been known for decades, most experiments are still performed at supraphysiological values, typically above 10 mM (86, 90, 102).

For beta cells to exert the coordinated function, a high level of cell-cell interactions within islets seem to be of utmost importance (53, 56, 71, 77, 83, 120, 141, 145, 147). Within areas of plasma membrane delimited by tight junctions, beta cells express gap junctions consisting of the connexin 36 protein that allows for electrical coupling and exchange of small signaling molecules between adjacent cells (35, 36, 103, 105, 131). Gap junctions, along with other intercellular signaling mechanisms, are thought to ensure coordinated cellular activity, narrowing the range of glucose concentrations stimulating insulin secretion at elevated glucose levels and preventing secretion in low glucose (18, 20, 28, 88, 133). However, the functional beta cell networks extracted from [Ca^2+^]_IC_ dynamics are much more heterogeneous than one would expect from a syncytium mediated by only gap junctions (14, 30, 34, 140). More specifically, instead of being relatively regular and lattice-like, the functional beta cell networks exhibit a heavy-tailed distribution, small-worldness, and a clustered structure with well-pronounced functional subcompartments (66, 84, 101, 106, 140).

As far as we know, the interplay between the abovementioned network parameters and characteristics of beta cell glucose responses is mostly unexplored, even more so when it comes to glucose-dependence of these relationships. Although the role of intercellular communication in islet dysfunction is incompletely understood, more and more studies suggest that the coordinated activity within islets is vital for normal insulin secretion dynamics and may be directly involved in glucose intolerance and the pathogenesis of diabetes mellitus (39, 78, 131, 141). Disruptions in gap junctional and paracrine communication abolish synchronized electrical and [Ca^2+^]_IC_ activity and lead to altered plasma insulin oscillations and to glucose intolerance (71, 116), as observed in different obesity and diabetes mellitus models (3, 18, 31, 43, 52, 53, 78, 137).

This study aimed to systematically measure and analyze, with high spatiotemporal resolution and over long periods of time, the dynamics of [Ca^2+^]_IC_ oscillations in many beta cells at a time with single-cell resolution in acute pancreas tissue slices to assess different glucose-dependent properties, with a particular emphasis on emergent collective operations in both physiological and supraphysiological glucose concentrations and their relationship with classical physiological properties. For this, we recorded beta cell activation, activity during the plateau phase, and deactivation, extracted and analyzed both classical and network functional parameters, and finally explored the relations between them. The latter is crucial for a deeper understanding of various aspects of recently reported heterogeneity within islets (8, 21, 55, 119, 122, 128, 129, 131).

## 2. Materials and Methods

### 2.1. Ethics statement

We conducted the study in strict accordance with all national and European recommendations on care and handling experimental animals, and all efforts were made to minimize the suffering of animals. The Administration of the Republic of Slovenia for Food Safety, Veterinary and Plant Protection approved the experimental protocol (permit number: U34401-12/2015/3).

### 2.2. Tissue slice preparation and dye loading

8-20 week old NMRI mice of either sex were kept on a 12:12 hours light: dark schedule in individually ventilated cages (Allentown LLC, USA) and used to prepare acute pancreas tissue slices, as described previously (136, 139). In brief, after sacrificing the mice, we accessed the abdominal cavity via laparotomy. We distally clamped the common bile duct at the major duodenal papilla. Proximally, we injected the low-melting-point 1.9 % agarose (Lonza, USA) dissolved in extracellular solution (ECS, consisting of (in mM) 125 NaCl, 26 NaHCO_3_, 6 glucose, 6 lactic acid, 3 myo-inositol, 2.5 KCl, 2 Na-pyruvate, 2 CaCl_2_, 1.25 NaH_2_PO_4_, 1 MgCl_2_, 0.5 ascorbic acid) at 40 °C into the common bile duct. Immediately after injection, we cooled the agarose infused pancreas with ice-cold ECS and extracted it. We prepared tissue slices with a thickness of 140 μm with a vibratome (VT 1000 S, Leica) and collected them in HEPES-buffered saline at RT (HBS, consisting of (in mM) 150 NaCl, 10 HEPES, 6 glucose, 5 KCl, 2 CaCl2, 1 MgCl2; titrated to pH=7.4 using 1 M NaOH). For staining, we incubated the slices for 50 minutes at RT in the dye-loading solution (6 μM Oregon Green 488 BAPTA-1 AM (OGB-1, Invitrogen), 0.03% Pluronic F-127 (w/v), and 0.12% dimethylsulphoxide (v/v) dissolved in HBS). All chemicals were obtained from Sigma-Aldrich (St. Louis, Missouri, USA) unless otherwise specified.

### 2.3. Stimulation protocol and [Ca^2+^]_IC_ imaging

We transferred individual tissue slices to a perifusion system containing 6 mM glucose in carbogenated ECS at 37 °C. We exposed them to a single square pulse-like stimulation per slice, with the following glucose concentration (in mM)/duration of stimulation (in minutes): 7/40, 8/30, 9/20, 12/15, or 16/15, followed by incubation in a solution with substimulatory 6 mM glucose concentration until all the activity switched off. We used concentrations typically encountered in normal mice *in vivo* (7, 8, and 9 mM) and two concentrations (12 and 16 mM) that lie within the range usually encountered in diabetic mice and are close to values typically used in previous studies (11.1 and 16.7 mM, corresponding to 200 and 300 mg/dl, respectively). The concentrations lie closer together near the threshold and further apart in the higher range for a good trade-off between resolution and range. Importantly, each islet was stimulated with a single stimulatory condition. A single stimulation pulse duration varied due to large differences in time needed to activate and deactivate beta cell networks at different glucose concentrations and to ensure a comparable number of [Ca^2+^]_IC_ oscillations that entered analyses from different glucose concentrations. To limit the required experiments and analyses to a practically manageable number, the relationships between classical and network parameters were analyzed for the 8 mM and 12 mM glucose protocols only. We performed the imaging on a Leica TCS SP5 AOBS Tandem II upright confocal system (20x HCX APO L water immersion objective, NA 1.0) and a Leica TCS SP5 DMI6000 CS inverted confocal system (20X HC PL APO water/oil immersion objective, NA 0.7). Acquisition frequency was initially set to 1-2 Hz at 512 x 512 pixels during the first phase response, allowing for the determination of response onsets and deactivation, and switched to 27 – 50 Hz at 128 x 128 pixels for an intermittent sampling of the plateau phase to allow for a more precise quantification of [Ca^2+^]_IC_ oscillations. Alternatively, a resolution of 10 Hz at 512 x 512 pixels was maintained throughout the whole stimulation protocol. OGB-1 was excited by a 488 nm argon laser, and the emitted fluorescence was detected by Leica HyD hybrid detector in the range of 500-700 nm (all from Leica Microsystems, Germany), as described previously (139).

### 2.4. Data analyses

We manually selected ROIs and exported traces for an off-line analysis utilizing a custom-made software application (ImageFiltering, copyright Denis Špelič). We excluded recordings with extensive motion artifacts. Further off-line analysis of [Ca^2+^]_IC_ traces were made using in-house MATLAB/Python scripts. Fluorescence signals were expressed as F/F0, the ratio of the fluorescence signal (F) at a particular time point of the experiment relative to the initial fluorescence (F_0_). To account for photobleaching, we used a combination of linear and exponential fitting. The methodology used to determine [Ca^2+^]_IC_ signal characteristics, e.g., the duration of oscillations established at a half-maximal amplitude of the spike, the number of oscillations per minute, and the percentage of active time, is described in detail in the respective figures and figure captions. The activation times and deactivation times, i.e., the start of [Ca^2+^]_IC_ increases/decreases after switching from basal to stimulatory concentration (and vice versa), were selected manually. We considered two cells to be in the same activation/deactivation cluster if their activation/deactivation times were less than 3 seconds apart. For the statistical analyses, we used SigmaPlot 11 and 14 (Systat Software Inc). Statistical differences between groups were tested using ANOVA on Ranks and posthoc Dunn’s method. Asterisks denote statistically significant differences (* p < 0.05, ** p < 0.01, *** p < 0.001).

### 2.5. Network analyses

To quantify the beta cell collective activity in each islet, we constructed functional connectivity networks. We considered two cells functionally connected if their activity profiles exceeded a preset degree of synchronization, as described elsewhere (66, 78, 140). The resulting functional networks were diagnosed with conventional network metrics. Specifically, we calculated the average degree and the relative degree distribution to explore the connectivity of cells. To evaluate the network’s traffic capacity and functional integration of individual cells, we computed the global efficiency and the largest component. To characterize the functional segregation, we calculated the clustering coefficient and modularity, which reflect the level of clique-like structures within interconnected cell assemblies and the extent of division into smaller subpopulations, respectively. For details, see (27, 65).

### 2.6. Comparing classical and network parameters

We examined whether there are any interrelations between different network parameters and classical physiological parameters extracted from the phases of activation, sustained activity, and deactivation. To that purpose, we divided the cells in each islet into three subgroups. One-sixth of the cells with the lowest and highest values were designated to the first and third group, respectively, and the remaining 2/3 of the cells with intermediate values to the second group. The cellular signaling characteristics that we compared were the response onsets in the activation phase, relative active time and node degree in the plateau phase, and cellular deactivation times after cessation of stimulation in the deactivation phase. To ensure direct comparability and pooling of the data from all islets, the active times and node degrees were normalized by dividing each value by the average value in the given islet. For example, if a cell had a normalized active time 1.05 and normalized node degree 0.9, its activity was 5 % higher than the average, whereas the number of connections was 10 % lower. For the activation and deactivation sequence, we used relative ranks. For the first and last responding cell, the activation and deactivation ranks were 0 and 1, respectively, irrespective of the number of cells or the absolute values of time in the given islet. We performed a separate analysis for islets stimulated with 8 mM and 12 mM glucose.

## 3. Results

The response of a typical beta cell consisted of three subsequent phases, i.e., activation, plateau, and deactivation. Below, we present their physiological and network properties in this logical order.

### 3.1 Glucose-dependence of beta cell activation

Beta cells responded to a given glucose concentration with a delay in the onset of [Ca^2+^]_IC_ increase (activation delay) that progressively shortened with increasing glucose concentrations (Figure 1, red). Strikingly, the median beta cell activation delay was more than 12 min (728 s) at the threshold concentration of 7 mM glucose and progressively decreased to about 1.5 min (104 s) at 16 mM glucose (Figure 1C). We observed a large degree of heterogeneity in delays among individual cells and cell clusters at a given glucose concentration. Interestingly, this heterogeneity was glucose-dependent, as evident from progressively shorter lags between the first-responding cell’s response in a given islet and any other cell’s response in this same islet (first cell-any cell delay; Figure 1, blue). More specifically, the interquartile range in first cell-any cell delays decreased from around 12 minutes (728 s) in 7 mM glucose to less than half a minute (17 s) in 16 mM glucose (Figure 1D). Consistent with the above findings, the time when 50 % of cells activated decreased progressively with increasing glucose concentrations (from 450 s at 7 mM to 17 s at 16 mM, Figure 1E).

**Figure 1:**
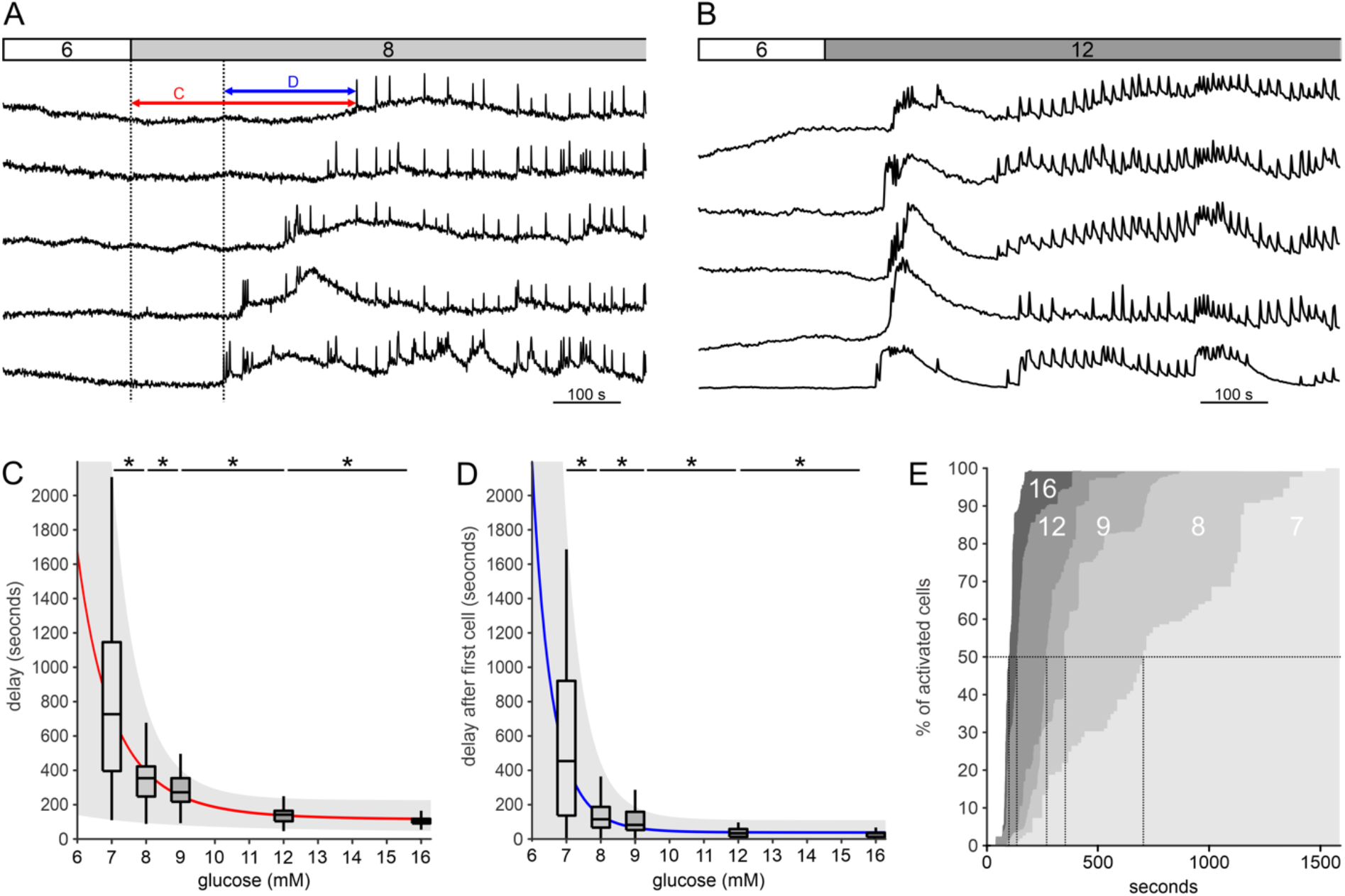
Glucose-dependent activation. **A-B** Response onsets of typical beta cells to stimulation with 8 mM (A) and 12 mM glucose (B). Lines indicate activation delays in individual cells (red line, pooled data shown in panel C) and variability of delays (blue line, pooled data shown in panel D). **C** Glucose-dependence of activation delays after stimulation with 7 mM, 8 mM, 9 mM, 12 mM, and 16 mM glucose. A power-law fit through median values (red line) and the inter-whiskers interval (gray area) are added for better visualization. 1st quartile/**median**/3^rd^ quartile (Q1/**M**/Q3), in seconds): 396/**728**/1148 (7 mM), 248/**355**/423 (8 mM), 217/**273**/355 (9 mM), 104/**142**/168 (12 mM), and 91/**105**/121 (16 mM). D Glucose-dependence of first cell-any cell delays. Q1/**M**/Q3 (in seconds): 130/**542**/920 (7 mM), 62/**112**/184 (8 mM), 48/**79**/155 (9 mM), 10/**26**/50 (12 mM), and 7/17/33 (16 mM). **E** Cumulative distributions of activation delays within islets. Vertical lines indicate the time at which half of the cells were activated at a given stimulus (in seconds): 450 (7 mM), 111(8 mM), 78 (9 mM), 25 (12 mM), and 17 (16 mM). Slopes of sigmoidal dose-reponse fit: 0,002 (7 mM), 0,004 (8 mM), 0,007 (9 mM), 0,017 (12 mM), and 0,023 (16 mM). Data pooled from the following number of cells/islets: 137/7 (7 mM), 410/10 (8 mM), 378/6 (9 mM), 371/12 (12 mM), 498/7 (16 mM).

As observed from Figure 1E, the activation of beta cells during stimulation had a staircase-like appearance, suggesting that at least at the temporal resolution used in this study, groups of beta cells, rather than single cells, activated at the same time. Closer inspection of this pattern in the recorded time series revealed that beta cells formed spatiotemporal clusters of neighboring cells in response to glucose stimulation (Figure 2). To quantify the relationship of cluster sizes to glucose concentration, we calculated the percentage of arbitrarily defined large clusters (> 33 %, > 50%, and > 67 % with respect to all cells in the islet) for different glucose concentrations (Figure 2E). The clusters of activated cells were larger in higher glucose concentrations, as demonstrated by the cluster sizes’ cumulative distribution for different glucose concentrations (Figure 2F).

**Figure 2:**
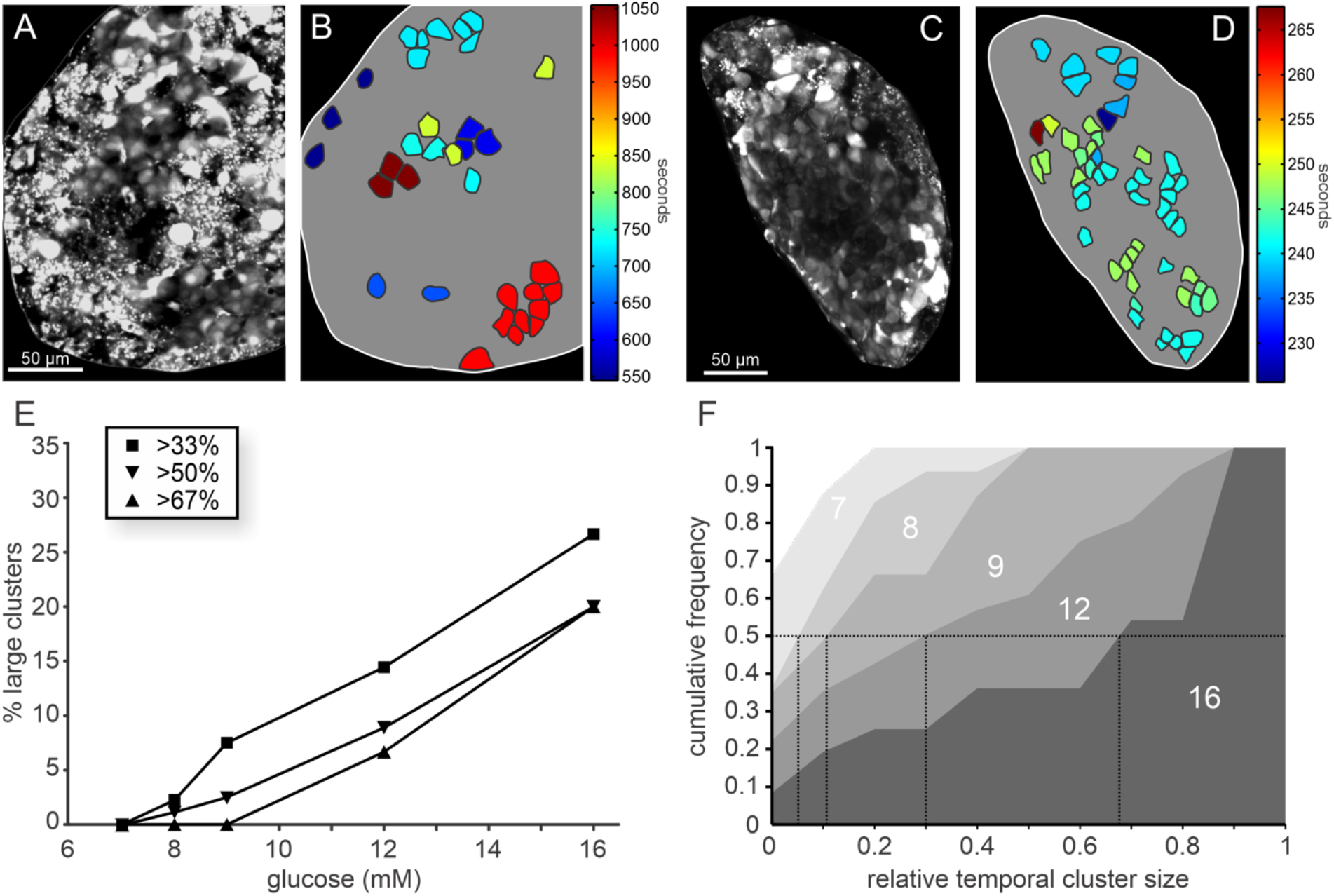
Spatiotemporal characterization of beta cell activation: **A-D** Color-coded response onset times of beta cells after 8 mM (B) and 12 mM (D) glucose stimulation. A and C show OGB-1 loaded cells in an islet. **E**: Distribution of relative sizes of clusters (relative to all active cells), signifying simultaneously activated cells under stimulation with different glucose concentrations. **F** Cumulative distribution of the relative cluster sizes for different glucose concentrations. Vertical lines indicate the relative temporal cluster size during activation of the first half of the cells in an islet: 0 (7mM), 0.05 (8 mM), 0.11 (9 mM), 0.30 (12 mM), and 0.68 (16 mM).

### 3.2 Glucose-dependent spatiotemporal [Ca^2+^]_IC_ dynamics during the plateau phase

#### 3.2.1 Classical functional parameters

Following activation, the plateau phase was characterized by relatively regular fast [Ca^2+^]_IC_ oscillations that, especially at higher stimulation levels, encompassed most cells in an islet. Increasing glucose concentrations characteristically affected both the frequency and duration of oscillations (Figure 3). More specifically, the frequency of oscillations increased across the physiological range of glucose concentrations (7-9 mM, Figure 3A), reached its peak at 12 mM, and decreased to 16 mM glucose (Figure 3 C). On the other hand, oscillation durations did not seem to be modulated at physiological glucose concentrations but progressively increased in 12 and 16 mM glucose (Figure 3D). As a consequence, the relative active time, i.e., the percentage of time that cells spend at an increased [Ca^2+^]_IC_, was dominated by the increasing frequency of oscillations in the range from 7 to 9 mM glucose, and by increasing duration of [Ca^2+^]_IC_ oscillations in higher concentrations. This resulted in a strikingly linear concentration-dependence of active time in the range between 7 and 16 mM glucose (Figure 3E), with the slope of the linear fit, *k* = 0.043/mM. This slope indicates that the active time increased by roughly 5 % for each 1 mM increment in glucose, starting from substimulatory 6 mM (AT=0), and achieved (by extrapolation) 100 % at 23 mM glucose. As explained further in Discussion, the fit is expected to become non-linear at higher glucose, as also suggested by previous studies, the abovementioned concentration at 100 % of active time is thus probably an underestimation (i.e., the sensitivity is overerstimated). As presented in boxplots, we noticed a great variability of both oscillation durations and frequencies at each glucose concentration. Notably, across different islets within each glucose concentration, there was a clear tendency of [Ca^2+^]_IC_ oscillations to last longer when their frequency was less and *vice versa*. Therefore, in Figure 3F, we provide an additional presentation of how active time depends on concentration, offering an even further insight into beta cells’ coding. More specifically, plotting the average oscillation duration as a function of the average inter-spike interval enables one to simultaneously represent three key parameters graphically: the frequency (i.e., the inverse value of the inter-spike interval), the duration, and the active time, represented here as the correlation slope of the two parameters, reflecting the ratio between duration and inter-oscillation interval of the given oscillation (in the given islet). This approach corroborates the finding that the active time is strongly concentration-dependent, since the slope (active time) increases with increasing glucose. At the same time, it quantitatively confirms the strong inverse relationship between duration and frequency observed across different islets at a given glucose concentration.

**Figure 3:**
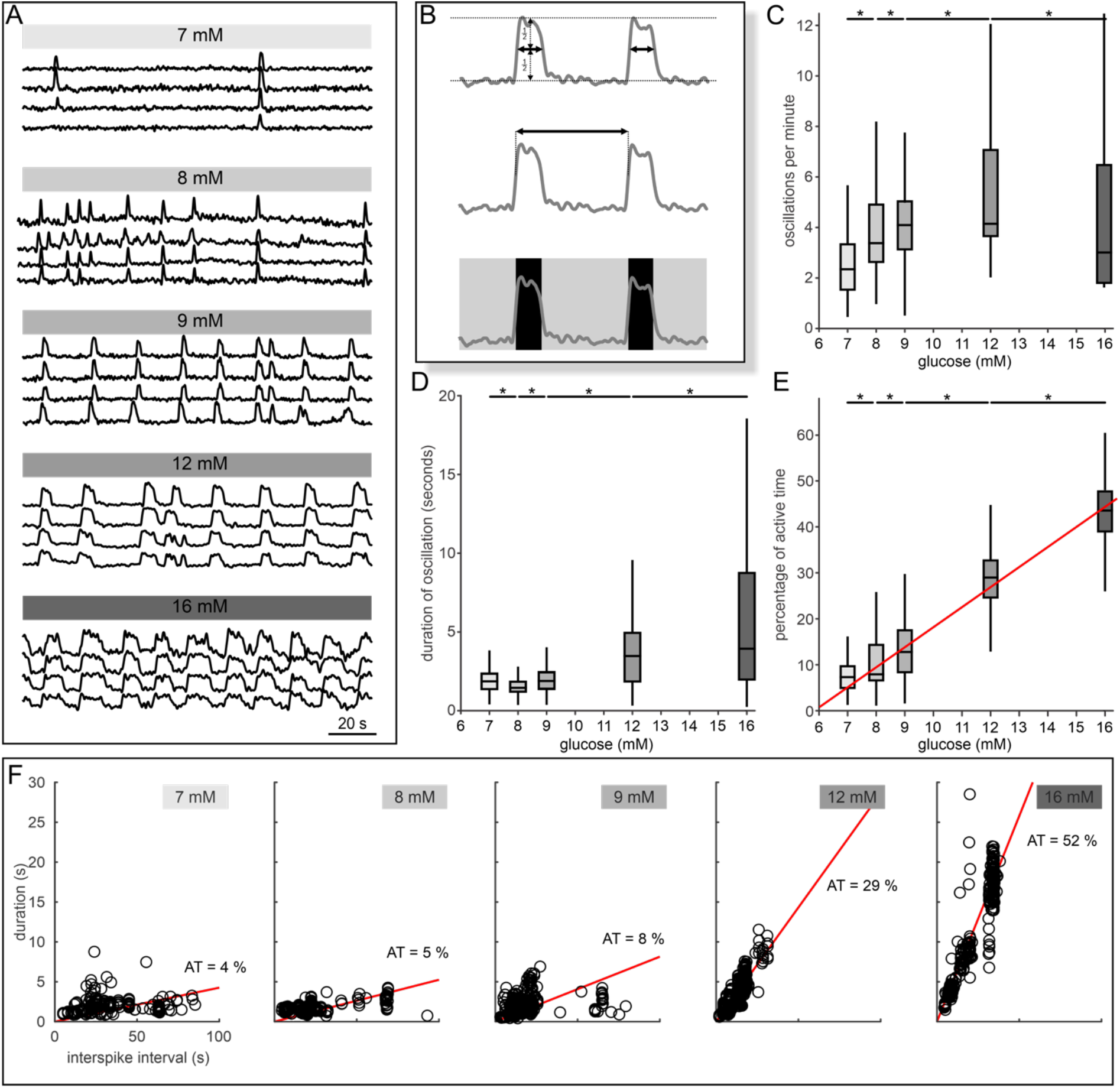
Spatiotemporal characterization of the plateau phase. **A** The oscillatory [Ca^2+^]_IC_ activity during stimulation with 7 mM, 8 mM, 9 mM, 12 mM, and 16 mM glucose. Shown are four representative cells from an islet per stimulus. **B** Schematic presentation of analyzed parameters: duration of oscillations at half-maximal amplitude (upper panel), number of oscillations per minute (middle panel), and percentage of active time (lower panel). **C** Frequency of oscillations. Q1/M/Q3 (in minute^−1^): 1,5/**2,4**/3,3 (7 mM), 2,6/**3,4**/4,9 (8 mM), 3,1/**4,1**/5,0 (9 mM), 3,7/**4,2**/7,1 (12 mM), and 1,8/**3,0**/6,5 (16 mM). **D** Duration of oscillations. Q1/M/Q3 (in seconds): 1,4/**1,8**/2,4 (7 mM), 1,2/**1,5**/1,9 (8 mM), 1,4/**1,9**/2,4 (9 mM), 1,9/**3,5**/4,9 (12 mM), and 2,0/**3,9**/8,7 (16 mM). E Percentage of active time. Q1/M/Q3 (in %): 5/**7**/9 (7 mM), 7/**8**/14 (8 mM), 8/**13**/18 (9 mM), 25/**29**/33 (12 mM), and 39/**44**/48 (16 mM). The red line indicates a linear fit through medians (R^2^=0.99). **F** Durations of individual oscillations as a function of the respective interspike interval. Glucose concentrations and slopes (designated as active time, AT) of linear regression lines are indicated. Data pooled from the following number of cells/islets: 171/8 (7mM), 241/7 (8 mM), 350/16 (9 mM), 392/16 (12 mM), and 281/6 (16 mM).

### 3.2.2 Network functional parameters

To further characterize beta cell collective behavior during the plateau phase, functional networks were constructed for each glucose concentration, as described in the Methods section. The results are presented in Figure 4. As shown previously, functional connectivity within an islet evolved with increasing stimulatory glucose concentrations (101). Stimulation with low glucose concentrations (7 or 8 mM) yielded mostly isolated and seldom synchronized beta cell activity. Increasing glucose resulted in greater coordination of cellular activity within an islet, demonstrated by the increasing density of networks (Figure 4A-E), greater average correlation in activity, and higher node degrees (Figure 4G and H). The increase in synchronization can be explained by increased activity, i.e., a greater number of [Ca^2+^]_IC_ oscillations and an increased number of cells involved in individual oscillations. The relative degree distributions for different glucose concentrations presented in Figure 4F show that beta cell networks are, in general, very heterogeneous. Except for very high glucose concentrations, a relatively small fraction of cells existed, which were very well connected. In 7 and 8 mM glucose, these cells were functionally correlated with up to 20-30 %, whereas in 9 and 12 mM glucose, they were correlated with up to 60 % of other beta cells. At 16 mM, the network became very dense, with most of the cells having a high number connections (Figure 4E and F). Intense stimulation diminished the intrinsic cellular variability, and the spatiotemporal activity was dominated by global and fully synchronized [Ca^2+^]_IC_ oscillations. Increasing glucose concentrations also resulted in functional networks that were more integrated, both locally, as illustrated by an increase in average clustering (Figure 4J), and globally, as demonstrated by decreased modularity (Figure 4K). Moreover, the rise in glucose concentration led to more robust networks with higher levels of functional integration, as evident from an increase in both the relative largest component and the global efficiency (Figures 4I and 4L).

**Figure 4:**
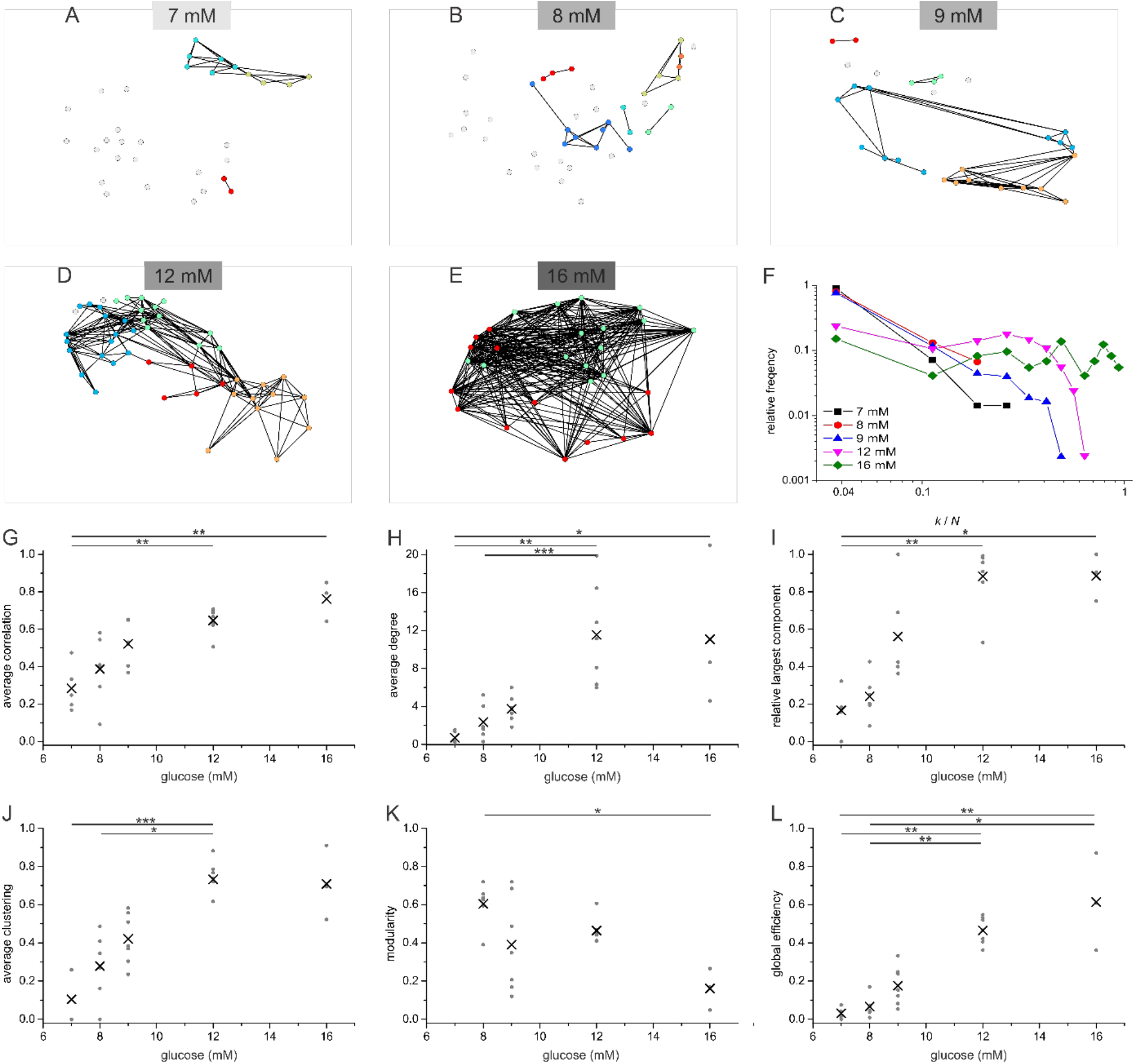
Beta cell functional connectivity at different glucose concentrations. **A-E** Characteristic functional networks for 7 (A), 8 (B), 9 (C), 12 (D), and 16 mM glucose (E). **F** Relative degree distributions at different glucose concentrations combined and normalized for all functional networks at a given stimulatory condition; k denotes node degree, and N denotes the number of all cells in a given islet. **G-L** Synchronization and network metrics as a function of glucose concentration: average correlation coefficient (panel G, median (M)/glucose concentration (GC, in mM): 0.25/7, 0.41/8, 0.52/9, 0.66/12, and 0.79/16), average network degree (panel H, M/GC (in mM): 0.33/7, 1.74/8, 3.27/9, 12.86/12, and 8.66/16), relative largest component (panel I, M/GC (in mM): 0.17/7, 0.23/8, 0.41/9, 0.91/12, and 0.90/16), average clustering coefficient (panel J, M/GC (in mM): 0/7, 0.31/8, 0.38/9, 0.72/12, and 0.70/16), modularity (panel K, M/GC (in mM): 0.25/7, 0.63/8, 0.35/9, 0.44/12, and 0.17/16; note that in 7 mM the networks were too sparse for a firm calculation of the modularity), and global efficiency (panel L, M/GC (in mM): 0.03/7, 0.06/8, 0.12/9, 0.43/12, and 0.61/16). Grey dots indicate values for individual islets and the black crosses the averages over all islets at a given concentration. Data pooled from the following number of cells/islets: 158/5 (7 mM), 343/6 (8 mM), 434/7 (9 mM), 413/9 (12 mM), and 73/3 (16 mM).

### 3.3 Glucose-dependence of beta cell deactivation

Removal of stimulatory glucose led to a concentration-dependent deactivation of beta cells (Figure 5, red). We observed a clear concentration-dependence for delays before the onset of deactivation (deactivation delay), which were less than 3 min (median delay 174 s) at 7 mM glucose. After stimulation by higher glucose concentrations, beta cells remained active longer. This delay reached a maximum value of about 15 min (median delay 815 s) after 16 mM (Figures 5C and E). The differences between cells of the same islet, as judged by first cell-any cell delays, also became larger with higher glucose (Figure 5, blue). However, this trend was less well pronounced than the inverse trend during activation. Additionally, for 7, 8, and 9 mM glucose, the differences between cells of the same islet were smaller during deactivation than during activation, whereas for 12 and especially 16 mM glucose, they became larger (compare first cell-any cell activation and deactivation delays in Figures 1 and 5).

**Figure 5:**
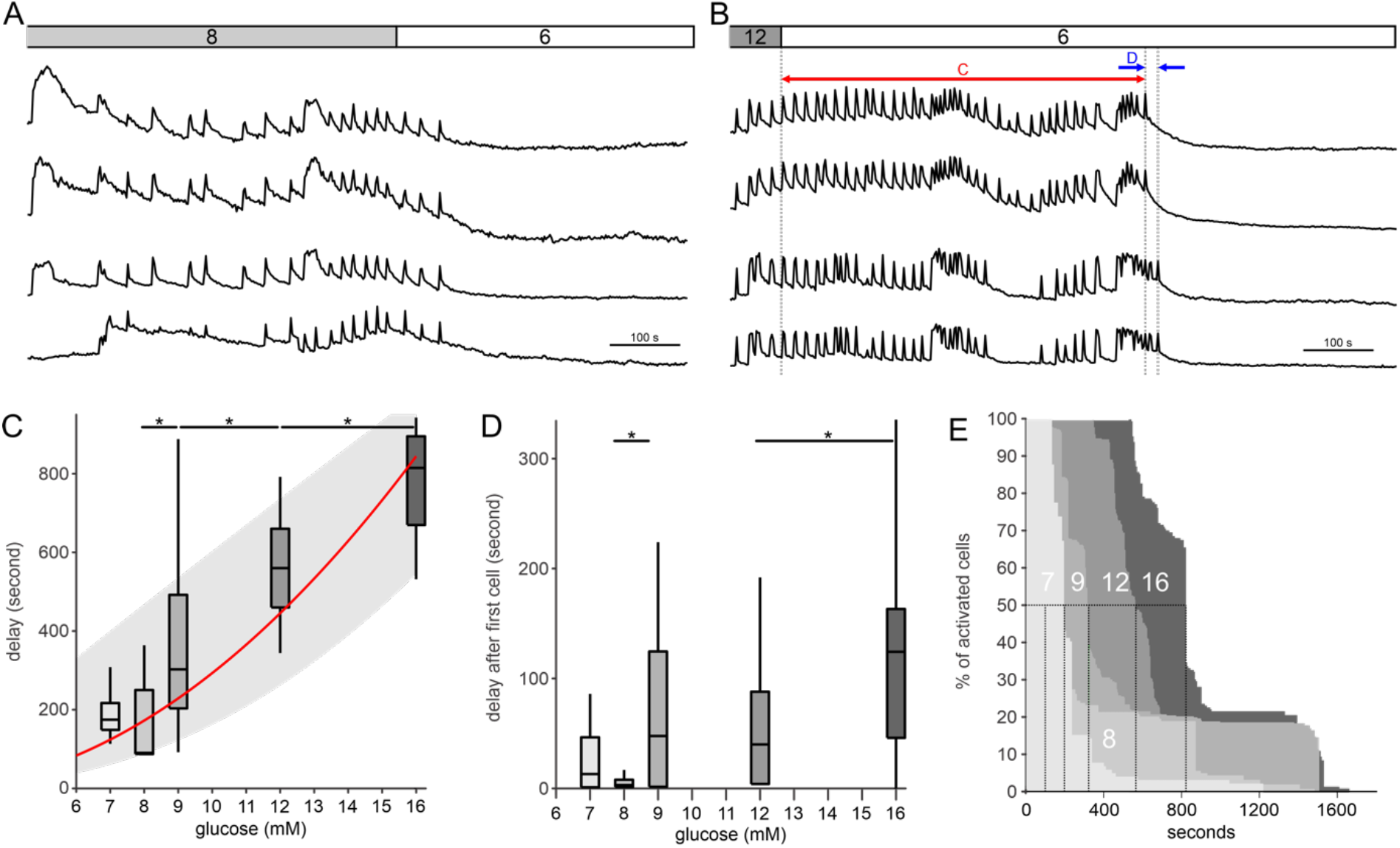
Glucose-dependent deactivation. **A-B** Deactivation of beta cells after cessation of stimulation with 8 mM (A) and 12 mM (B) glucose. Lines indicate deactivation delays in individual cells (red, pooled data are shown in panel C) and any-cell-first-cell deactivation delays (blue, pooled data shown in panel D). **C** Deactivation delays after stimulation with 7 mM, 8 mM, 9 mM, 12 mM, and 16 mM glucose. A power-law fit through median values (red line) and through the inter-whisker intervals (gray area) are added for better visualization. Q1/**M**/Q3: 149/**175**/217 (7 mM), 87/**90**/250 (8 mM), 203/**303**/492 (9 mM), 460/**560**/660 (12 mM), and 670/**815**/895 (16 mM). D Glucose-dependence of first cell-any cell deactivation delays. Q1/**M**/Q3: 1/**13**/48 (7 mM), 1/**3**/8 (8 mM), 1/**48**/125 (9 mM), 4/**40**/88 (12 mM), and 46/**124**/163 (16 mM). **E** Cumulative distributions of delays between the end of stimulation and deactivated at a given stimulus (in seconds): 193 (7 mM), 100 (8 mM), 316 (9 mM), 560 (12 mM), and 816 (16 mM). Data pooled from the following number of cells/islets: 107/7 (7 mM), 164/5 (8 mM), 372/6 (9 mM), 366/12 (12 mM), 197/6 (16 mM).

As with activation, deactivation occurred in spatiotemporal clusters (Figure 6), a phenomenon observed at both physiological (Figure 6A-B) and supraphysiological glucose concentrations (Figures 6C-D). The abovementioned increase in intercellular heterogeneity in higher glucose was evident in the decrease in cluster size of cells that deactivated together in higher glucose concentrations. The sizes of these clusters decreased with increasing glucose concentrations (Figures 6E-F). This strongly contrasts with the activation properties where cluster sizes were larger in higher glucose concentrations (compare Figures 2 and 6).

**Figure 6:**
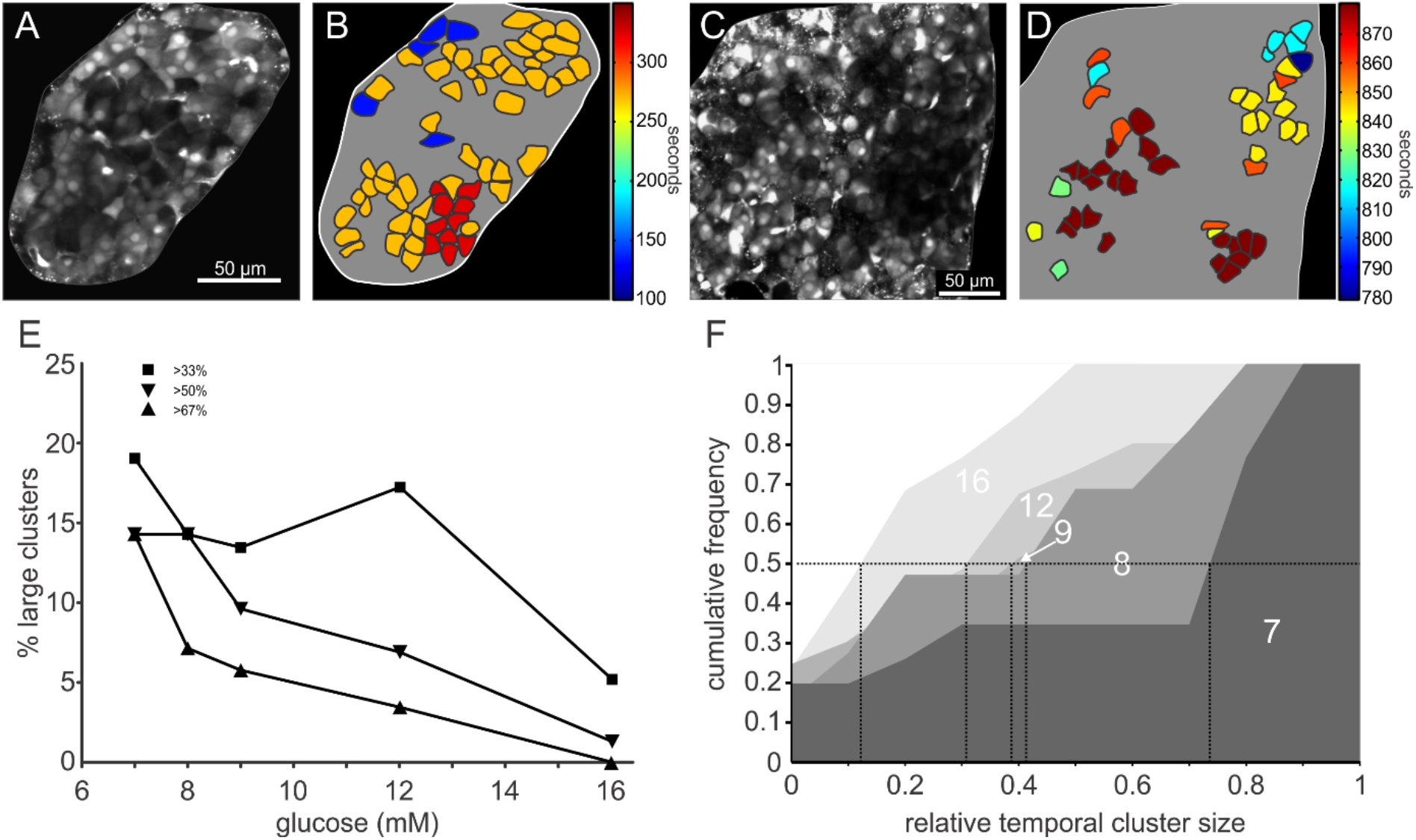
Spatio-temporal characterization of beta cell deactivation. **A-D** Color-coded response deactivation after cessation of stimulation with 8 mM (B) and 12 mM (D) glucose. A and C show the loading of cells with OGB-1 for respective islets. **E** Distribution of relative sizes of clusters, signifying simultaneously deactivated cells after removal of stimulus. **F** Cumulative distribution of relative cluster sizes. Vertical lines indicate the relative temporal cluster size during deactivation of the first half of the beta cells in an islet.

### 3.4 Comparison between activation, plateau activity, and deactivation

We compared the three phases of beta cell response in Figure 7. Figures 7B-D show how the one sixth of cells with shortest delays before activation (i.e., the first-responders) compare with the two third majority of cells with average delays and the one sixth of cells with the longest delays (i.e., the last-responders), with respect to active time and node degree during the plateau phase, as well as the deactivation time rank, for two selected concentrations of glucose, i.e., 8 and 12 mM. The first-responders were the most active group in the plateau phase in 12 mM glucose and more active than the last responders in 8 mM glucose (Figure 7B). The first-responders also had the most functional connections in 12 mM glucose and more than the last-responders in 8 mM glucose (Figure 7C). Surprisingly, the relation with deactivation seemed to be glucose-dependent, with the first-responders being the last to stop their activity after exposure to 8 mM glucose and the last-reponders being the last to stop their activity after exposure to 12 mM glucose (Figure 7D). The relationship between both parameters calculated during the plateau, i.e., the active time and the node degree is displayed in Figure 7E. The cells with the most functional connections, i.e., hubs, had longer active times than the cells with the least functional connections in 8 mM glucose, whereas there were no significant differences in 12 mM glucose. Furthermore, in both glucose concentrations the cells that deactivated the first (i.e., the first-deactivators) had the shortest active times (Figure 7F). Finally, the first-deactivators had the least functional connections in 8 mM glucose and less connections than the majority in 12 mM glucose (Figure 7G). The remaining 6 pairwise comparisons (i.e., activation time rank, deactivation time rank, and node degree vs. relative active time; activation time rank vs. deactivation time rank; activation time rank and deactivation time rank vs. node degree) are shown in Figure S1)

**Figure 7:**
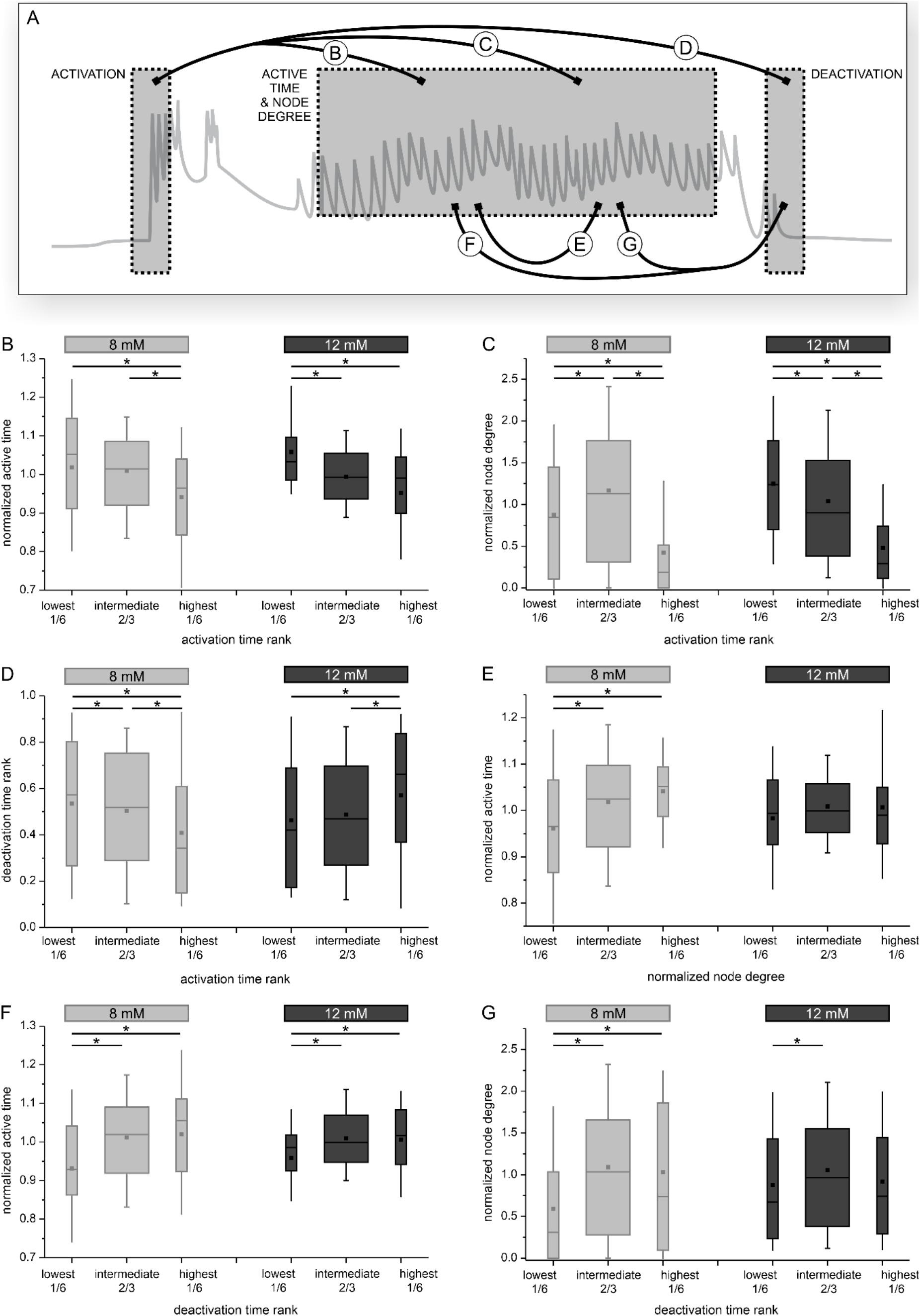
The relationship between functional parameters during activation, plateau activity, and deactivation, in 8 mM (light grey) and 12 mM (dark grey) glucose. **A** The analyzed parameters with respect to the phases of a typical response. Letters indicate panels with pairwise comparisons. **B-D** The relationship between response onset (activation rank) and the relative active time (B), node degree (C), and deactivation rank (D). **E** The relationship between node degree and relative active time during the sustained plateau activity. **F-G** The relationship between deactivation time rank and active time (F), as well as node degree (G). The analysis was performed on a particular set of islets subjected to sustained and long-lasting recordings (~1 hour). Data pooled from the following number of cells/islets: 587/7 (8 mM) and 860/9 (12 mM).

## 4. Discussion

For clarity, we discuss the importance of our findings in the logical order that follows a typical beta cell response to glucose and which was already employed above, i.e., from activation to deactivation.

### 4.1. Activation

We wish to first addres the possibility that the observed behavior, at least during activation and deactivation, is an experimental artifact due to dynamics of glucose increases in our setup in various parts of the islet. However, as we demonstrated before by using a large extracellular dye to quantify perifusion in our chamber, we believe that this is most certainly not the case and that beta cells are quickly exposed to practically identical extracellular glucose concentrations (139). In previous studies, the activation appeared either synchronous among cells (26) or varied to some degree (44, 59, 62, 78, 81, 85, 99). In either case, to the best of our knowledge, this property has not been systematically analyzed nor emphasized, with the exception of our previous reports using in total only two different concentrations of glucose (8 and 12 mM) (139, 142). The present study confirmed our previous findings for these two particular concentrations, using an independent large set of cells and islets from different animals. More importantly, here we employed a whole range of concentrations and beta cell activation profiles exhibited pronounced glucose-dependence and considerable variability in time and space (Figure 1). More specifically, we detected a median response delay of 15 minutes at the threshold of 7 mM glucose, which became progressively shorter and reached a median delay of 2 minutes at the highest tested concentration of 16 mM glucose. We propose this phenomenon be termed “advancement” of beta cell activation. A similar temporal profile of metabolic activity observed after stimulation with different glucose concentrations directly corroborates the advancement observed in this study, suggesting that the time needed for the cells to metabolize enough glucose to become active becomes progressively shorter with increasing stimulus intensity (48, 81). Physiologically, an earlier response to a stronger stimulus may play an important role in preventing too large excursions of glucose *in vivo*. Interestingly, during the first couple of minutes following activation and before a sustained pleateau phase was established, an exceedingly high frequency of fast oscillations or a [Ca^2+^]_IC_ signal that could be explained by continuous bursting of electrical activity were observed, indicating that during this time, there is a transient “overshoot” of activity, as observed before (59, 91, 111, 117, 139, 142). Further studies are needed for a more detailed quantitative description and to determine the mechanistic substrate of this phenomenon, but from a homeostatic perspective, this behavior could indicate that beta cells behave as both phasic and tonic sensors, responding differently to an increasing and a constantly increased glucose, again ameliorating excursions of glycemia *in vivo (89, 98, 100, 107)*. This is especially important considering that *in vivo*, [Ca^2+^]_IC_ may actually follow an oscillatory rather than a sustained profile (42, 67, 109), that [Ca^2+^]_IC_ dynamics can largely explain biphasic insulin secretion (108, 111), and that the loss of first-phase insulin secretion is an early sign of T2DM (32, 115).

Moreover, we demonstrated large heterogeneity between individual cells within the same islet: at the threshold glucose level, the first cells responded after ~3 minutes, and the last cells more than 25 minutes later. This variability indicates large differences in their sensitivity to glucose. Due to a limited number of concentrations and time periods employed in our study, we cannot exclude the possibility that the actual threshold for activation actually lies a bit below 7 mM glucose, but at least in the NMRI strain and for our study design, cells never exhibited any activity at 6 mM glucose. However, since beta cells may also act as differential and not only difference sensors, we do not exclude the possibility that at least some cells may at least transiently respond to 6 mM glucose, in case they were exposed to lower concentrations before (e.g., 3 mM).

From a functional point of view, differences in sensitivity to glucose enable recruitment. Looking at the slope of the cumulative distributions in Figure 1, there was practically no cooperativity during recruitment at 7 mM glucose, strongly suggesting that at this concentration, active cells did not seem to appreciably increase the probability of inactive cells becoming active. In higher glucose concentrations, differences between cells did not just become shorter due to higher glucose and faster responses, but there seemed to be an important contribution of active cells increasing the probability of inactive cells to become active. This is probably a consequence of increasing length constants for the electrotonic spread of gap junctional currents due to increasing input resistance and increasing junctional conductance, of intercellular diffusion of other messengers or intermediates, or a combination of all. This temporal aspect of recruitment is further complemented and the proposed mechanism corroborated by the finding that at the employed temporal resolution, clusters of nearby cells were recruited together and these clusters became larger in higher glucose.

We do not wish to suggest that the first-responders are in any way necessary for islets to respond to glucose and that their role is fixed in time. It is reasonable to speculate that the removal or dysfunction of first-responders would make other cells become first-responders (49, 82, 122, 128, 129), and recent experimental data corroborated this (91). Even without removal or dysfunction, the metabolic status and physiological features change as beta cells mature (12). This suggests that a cell’s role may also dynamically change with time, possibly within hours. To address this possibility, repetitive stimulations over longer periods are required in the future. Finally, activation seems to be an attractive additional parameter for testing the effects of various drugs, but appropriately long experimental protocols are needed to account for the large heterogeneity between cells, especially if concentrations < 9 mM glucose are used.

### 4.2. Plateau activity

#### 4.2.1 Fast [Ca^2+^]_IC_ oscillations

A hallmark of the plateau phase during sustained stimulation with glucose are repetitive fast [Ca^2+^]_IC_ oscillations that correspond to bursts of membrane potential depolarization proximally and pulses of insulin secretion distally in stimulus-secretion coupling cascade (7, 13, 19, 23, 25, 59, 60, 125, 139). A few studies attempted to decipher how glucose modulates the properties of these fast [Ca^2+^]_IC_ oscillations, yielding somewhat contradictory results. For instance, a recent microelectrode array study demonstrated that within the physiological concentration range glucose increased only frequency (95). In contrast, sharp electrode and [Ca^2+^]_IC_ imaging studies showed that supraphysiological concentrations affected mostly oscillation duration, with little effect on the frequency of oscillations (7, 13, 76, 104). On the other hand, Cook’s research suggests that frequency is the crucial regulated parameter across a wide range of glucose concentrations (37, 38). In the present paper, we found that the dose-response relationship showed two different regimes (Figure 3). First, the frequency of oscillations followed the increase in glucose concentration within the physiological limits of glucose concentrations,, increasing from approximately 2/min to 4/min from 7 to 9 mM glucose, whereas in this range, the oscillation durations did not seem to change to a biologically relevant extent and remained close to 2 seconds long. At higher glucose concentrations, an increase in oscillation duration rather than in frequency became the predominant strategy of increasing activity. From 9 mM to 16 mM glucose, the oscillations increased in duration approximately two-fold, whereas the frequency showed an inverse U-shape behavior, reaching a peak at 9 and 12 mM glucose and decreasing at the highest concentration tested. Although we did not test concentrations beyond 16 mM, our findings are consistent with the view that ultimately, fast [Ca^2+^]_IC_ oscillations disappear due to continuous bursting at very high levels of stimulation. As a side note, with current [Ca^2+^]_IC_ imaging approaches, it is impossible to capture very fast [Ca^2+^]_IC_ oscillations that correspond to individual action potentials (47).

As a result, there was a strikingly linear increase in the active time over the range of glucose concentrations tested (Figure 3), a feature also observed in studies utilizing microdissected islets (38, 76). In our hands, a single beta cell spent less than 10 % of the time in an active state at the threshold concentration and the active time increased by approx. 5 % per each mM of glucose increase. At this rate, the oscillatory behavior would reach a sustained active state (i.e., an active time of 100 %) at approx. 25 mM glucose or somewhat higher, given that the dose-response curve is probably flatter at higher concentrations (124). This is in good agreement with previous insulin secretion data, as well as [Ca^2+^]_IC_ imaging and electrophysiological studies (9, 20, 41, 57, 74, 97). A theoretically important aspect of the dual behaviour of the concentration-coding principle is that a beta cell cannot “know” or sense the glucose concentration based on either the frequency or duration of oscillations alone. This is further supported by the finding that at every given glucose concentration, there was a wide range of oscillation durations and frequencies (and a strong inverse relationship between the two). In other words, at a given glucose concentration, some islets responded with longer and less frequent oscillations and others with shorter and more frequent oscillations. Recent research suggests that at least within an islet, a higher intrinsic frequency is observed in metabolically less active cells (146). It remains to be investigated whether islets with higher frequencies tend to be less metabolically active compared with other islets. However, the active time or the product of duration and frequency of oscillations, is the physiological parameter that very closely reflects the concentration of glucose which beta cells are exposed to. In turn, the active time is closely related to insulin secretion (6, 13, 22, 23, 61, 72, 126). A practically very important repercussion of the above is that when studying the add-on effects of physiological and pharmacological secretagogues, one should always keep in mind the stimulatory glucose concentration used and not focus on either the frequency or duration of oscillations alone, but always calculate the active time.

The relationship between [Ca^2+^]_IC_ oscillations and insulin release (9, 20, 41, 57, 74, 97) has recently been integrated into the so-called metronome model. This model predicts that an increase in glucose increases the active time of bursting electrical acitivity and fast [Ca^2+^]_IC_ oscillations, leaving the frequency of the underlying slow oscillations unaffected, and ultimately leads to an increase in the amplitude of insulin pulses (127). Importantly, the dynamics of insulin release cannot be entirely explained by membrane potential and [Ca^2+^]_IC_ dynamics, but probably also involve changes in mobilization and priming of insulin granules (111).

### 4.2.2 Functional networks

Beta cells within a mouse islet associate into a single large syncytium. More specifically, gap junctional coupling via connexin 36, as well as many additional cell-cell contacts, paracrine, and neuronal signaling mechanisms form the structural basis for complex interactions between cells (5, 45, 51, 92, 121, 144). These interactions enable beta cells to function more efficiently than a comparable number of dissociated cells, by partially overcoming their heterogeneity (112–114). However, some heterogeneity persists in beta cells in islets and recent research suggests that it plays a crucial role for normal beta cell function (21, 106, 118, 131). One of the important tools to quantify heterogeneity are classical physiological and network analyses of complex spatiotemporal [Ca^2+^]_IC_ dynamics (78, 140), which enable identification of cells that play important roles in an islet’s response to secretagogues, e.g., cells that respond first (i.e., first-responders), cells that start the [Ca^2+^]_IC_ waves that synchronize fast [Ca^2+^]_IC_ oscillations (i.e., pacemakers), cells that functionally interact with the most other cells (i.e., hubs), etc (19, 66, 78, 139, 140, 143, 146). These roles are probably crucial for generation of coordinated rhythmic activity (15, 18, 36, 66, 78, 84, 101, 120, 123, 140, 142) and there is growing evidence that both environmental and genetic factors may target connectivity and heterogeneity in the pathogenesis of diabetes mellitus (4, 53, 78, 131, 141, 145).

In the present study, we advanced our understanding of beta cell networks by confirming the glucose dependency of intercellular connectivity and by systematically comparing the obtained network properties with classical physiological characteristics at the level of individual cells. The latter aspect is covered in the last chapter. With respect to the first, we found that the functional networks are segmented, locally clustered, and heterogeneous for physiological glucose concentrations. In other words, from a functional point of view, this might be regarded as a concept of islets within islets. Higher stimulation led to a more synchronized behavior and hence to denser, more integral and efficient networks, functioning as a large single network. This importantly corroborates our previous findings, but on a larger dataset, on a wider range of glucose concentrations, and based on a different stimulation paradigm (66, 101). The above qualitative changes in functional connectivity patterns go hand in hand with a different nature of [Ca^2+^]_IC_ oscillations. Under low stimulatory conditions, on average the oscillations are present in a smaller fraction of cells, and they are rarer and more erratic. At glucose concentrations beyond 12 mM, [Ca^2+^]_IC_ oscillations become more frequent, global, and coherent, resulting in functionally more connected networks (66, 142). Our findings seem to suggest that in future studies aimed at assessing functional heterogeneity under normal and pathological conditions, it seems to be advisable to use lower stimulatory concentrations of glucose than typically used in the literature, i.e., up to 12 mM. Beyond this level, many functional properties seem to saturate and differences between cells become much less well pronounced. It should be noted that in the present work, we did not analyze in detail the properties of the [Ca^2+^]_IC_ waves that are the synchronizing mechanism coordinating fast [Ca^2+^]_IC_ oscillations (19, 47, 125, 143). Such an analysis would enable us to also study the properties of the cells that initiate fast [Ca^2+^]_IC_ oscillations, i.e., pacemakers. However, such an endeavour is technically and analytically more challenging and will be covered in a separate article.

### 4.3. Deactivation

Afer the end of stimulation, beta cells gradually suppressed their oscillatory activity, and their [Ca^2+^]_IC_ returned to the basal level. The spatiotemporal pattern of this “off” response showed clear glucose-dependence (Figure 5). In brief, after exposure to lower levels of glucose, the beta cells deactivated sooner and more homogeneously, i.e., with shorter first cell-any cell delays and in larger clusters than after exposure to higher glucose concentrations. To explain the observed behavior with respect to beta cell metabolism and intercellular coupling, further modelling and experimental studies are warranted. That being said, to our knowledge this is the first systematic and quantitative report on the glucose-dependence of beta cell deactivation and provides a reference point for future studies. It is reasonable to speculate that the longer delays and more heterogenous “off” responses in higher glucose are a direct consequence of a greater metabolic activation during stimulation and thus longer time required for the ATP production to fall below the stimulatory level. Noteworthy, changes in deactivation properties are present in Cx36 knockout mice, together with an increased basal and lower stimulated insulin secretion (71, 116, 133) and in a genetic model of the metabolic syndrome, together with islet hypertrophy, changes in intercellular coupling, and insulin hypersecretion (39). This indicates that changes in deactivation, similarly to changes in activation and plateau activity could be an early marker of beta cell functional adaptation and dysfunction during development of diabetes (63, 81, 141). Moreover, under normal conditions insulin and glucose levels oscillate in different species with a period on the order of magnitude of 10 minutes (64, 71, 94) and the loss of these oscillations is an early marker of diabetes (93). Our current and previous research suggests that the loss of oscillations could at least partly be due to hyperglycemia *per se*, in the sense that beta cells become unable to turn off soon enough to allow for periods of silence between subsequent pulses (66).

### 4.4. Connecting the dots: relations between activation, plateau, and deactivation

The temporal and spatial resolution in our current study, together with the large number of analyzed cells, gave us the opportunity to systematically look for the first time for a possible overlap between a cell’s role during activation, plateau, and deactivation. More specifically, with respect to every parameter size, cells were divided into six parts, with the middle two sixths being pooled together into a majority with medium parameter values. In other words, our first-responders, most active cells, hubs, and first-deactivators are cells that belonged to the one sixth (or roughly 17 %) of cells with the smallest relative activation ranks, highest relative active times, most links in functional networks, and smallest relative deactivation ranks. Importantly, we wish to point out that paramater values were rather continuous, with no clear extremes. Also, we do not believe or wish to suggest that removing the cells with the highest parameter values would dramatically alter the collective behavior (49). Moreover, it remains to be investigated whether these roles are relatively constant in time or change dynamically (80). Finally, due to some recent confusion in the literature with respect to nomenclature, it shall be pointed out that the name “first-responders” pertains to the beginning of the transient activation, and that the name “pacemakers” shall be reserved for the cells that start the fast plateau [Ca^2+^]_IC_ oscillations. As mentioned previously, due to technical reasons, the latter will be analyzed in a separate study.

All things considered, the first-responders tended to be among the most active cells during the plateau (and *vice versa*, the most active cells tended to have the shortest delays to activation; Figures 7B and S1A). The most active cells, in turn, tended to have longer delays to deactivation (and *vice versa*, the last-deactivators tended to have longer active times, Figures S1B and 7F). Further, the functionally most connected cells, i.e., the hubs, tended to have higher active times (in 8 mM glucose, with no differences in 12 mM glucose, Figure 7E), activate among the first (in 12 mM glucose and not among the last in 8 mM glucose, Figure S1E), and deactivate among the last (in 8 mM glucose, with no differences in 12 mM glucose, Figure S1F). There was no clear-cut direct relationship between activation and deactivation properties (Figures 7D and S1D). Moreover, in addition to the abovementioned relationship between most connected and most active cells, it should be pointed out that the least active cells had the least functional connections (in 8 mM glucose and less than the majority in 12 mM glucose, Figure S1C). With regard to the relationship between most connected cells and activation, it can be added that first responders had the highest node degrees in 12 mM glucose and higher node degrees than the last responders in 8 mM glucose (Figure 7C). Finally, with regard to the relationship between most connected cells and deactivation, it seems that the first-deactivators are not among the most connected cells (Figure 7G).

Conceivably, a cell’s ability to activate soon, maintain high levels of activity and functional connections during the plateau phase, and turn off late, compared with other cells, may have to do with its relative higher metabolic activity and rates of ATP production, relatively smaller KATP conductance, which would enable it to reach the threshold for bursting activity sooner, and strong coupling with similarly excitable cells or weak coupling with much less excitable cells, which would not clamp it at resting membrane potential values too strongly, but help it activate and maintain active for longer. Validation of these hypotheses will require further testing by models and simultaneous or serial experimental assessment of metabolic and electrophysiological parameters and is beyond the reach of our current study. That said, we find it encouraging that our findings are compatible with previous modelling, electrophysiological and imaging studies, and may even help reconciliate some recent opposing views (49, 96, 119, 122, 128, 129). More specifically, different lines of evidence suggest that the functionally most connected cells are metabolically more active (21, 68, 84, 96, 122, 123). At the same time, our current and previous experimental and model findings that the parameter value distributions and functional behavior of cells are heteregoneous in a rather continuous manner (67, 68, 87, 101, 139, 140, 142), with no clear-cut subpopulations, are compatible with the view that only a few cells with extreme parameter values are not necessary for normal beta cell activation, coordinated activity, and deactivation (49, 129). Again, we would like to underline that the terms “first-responders” and “hubs”, as defined in this study, are not the same as “pacemakers”. We believe that the term pacemakers shall be strictly used for cells initating well-defined plateau fast [Ca^2+^]_IC_ oscillations. Our preliminary analyses suggest that the pacemaker role does not necessarily overlap with first-responders and hubs, which is again compatible with the view that pacemakers could be the metabolically least active cells (49).

In sum, we believe that beta cell heterogeneity with respect to activation, activity, and inactivation should become indisputable and that great strides shall be made in the future to better define terminology in this fascinating subfield of beta cell research, bring closer the researchers with different expertise and differing views on this matter, and to mechanistically explain the heterogeneity and define its clinical relevance.

## 4.1. Acknowledgements

We thank Maruša Rošer, Rudi Mlakar, and Maša Čater for their excellent technical assistance.

## 4.4. Author Contributions

### Conceptualization

Andraž Stožer, Marko Gosak, Marjan Slak Rupnik, Jurij Dolenšek

### Formal analysis

Andraž Stožer, Maša Skelin Klemen, Marko Gosak, Lidija Križančić-Bombek, Viljem Pohorec, Jurij Dolenšek

### Software

Jurij Dolenšek, Marko Gosak

### Experimental work

Andraž Stožer, Maša Skelin Klemen, Lidija Križančić-Bombek, Viljem Pohorec, Jurij Dolenšek

### Funding acquisition

Andraž Stožer, Marjan Slak Rupnik

### Project administration

Andraž Stožer, Marjan Slak Rupnik, Jurij Dolenšek

### Supervision

Andraž Stožer, Jurij Dolenšek

### Visualization

Andraž Stožer, Marko Gosak, Jurij Dolenšek

### Writing – original draft

Andraž Stožer, Maša Skelin Klemen, Marko Gosak, Lidija Križančić-Bombek, Viljem Pohorec Marjan Slak Rupnik, Jurij Dolenšek

### Writing – review & editing

Andraž Stožer, Marjan Slak Rupnik, Jurij Dolenšek

