## Supplemental Figure 1 for "Glucose-dependent activation, activity, and deactivation of beta cell networks in acute mouse pancreas tissue slices"

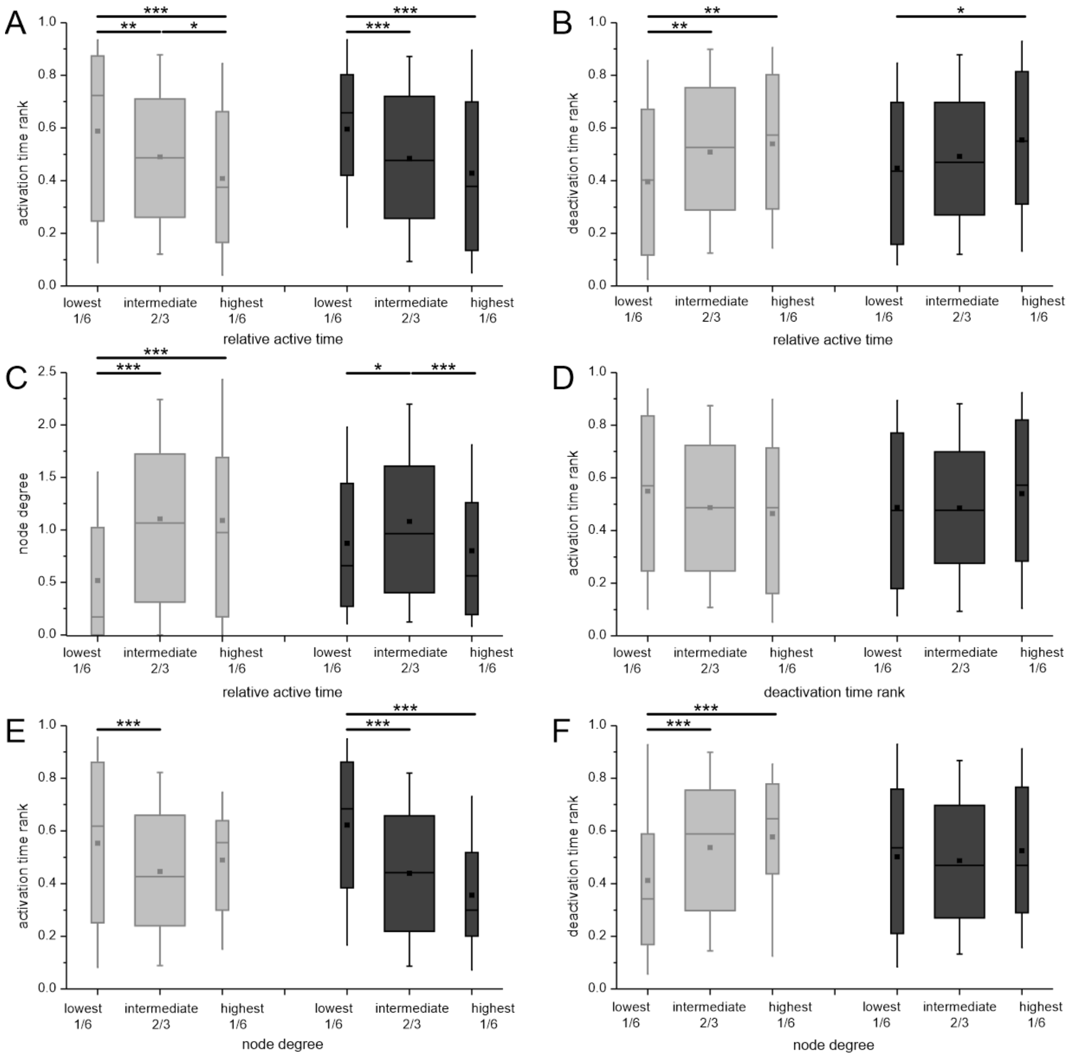


**Supplementary figure 1: The relationship between functional parameters during activation, plateau activity, and deactivation, in 8 mM (light grey) and 12 mM (dark grey) glucose. A-C:** The relationship between the relative active time and **(A)** 1$^{st}$ phase response onset (activation time rank), (**B**) deactivation of activity following stimulus withdrawal (deactivation time rank), and (**C**) normalized node degree. **D:** The relationship between activation and deactivation time ranks. **E-F:** The relationship between node degrees and (**E**) activation and (**F**) deactivation time ranks. The analysis was performed on a particular set of islets subjected to sustained and long-lasting recordings (~1 hour). Data pooled from the following number of cells/islets: 587/7 (8 mM) and 860/9 (12 mM). Note that this figure represents inverse relationships compared with Figure 7 in the main manuscript.
